# Connectomic analysis reveals an interneuron with an integral role in the retinal circuit for night vision

**DOI:** 10.1101/2020.02.24.963868

**Authors:** Silvia J.H. Park, Evan M. Lieberman, Jiang-Bin Ke, Nao Rho, Padideh Ghorbani, Pouyan Rahmani, Na Young Jun, Hae-Lim Lee, In-Jung Kim, Kevin L. Briggman, Jonathan B. Demb, Joshua H. Singer

**Author notes:** denotes equal contributions (Silvia J.H. Park, Evan M. Lieberman). **Author Contributions:** Conceptualization, JBD and JHS; Methodology, KLB; Software, KLB and NR; Formal Analysis, ELM and SP; Investigation, PG, NYJ, ELM, PR, and SP; Resources, HL, KLB; Writing – Original Draft, Review, and Editing JBD and JHS; Supervision, JBD, I-JK, JHS. **Declaration of Interests:** The authors declare no competing interests.

## Abstract

The mammalian rod bipolar (RB) cell pathway is perhaps the best-studied circuit in the vertebrate retina. Its synaptic interactions with other retinal circuits, however, remain unresolved. Here, we combined anatomical and physiological analyses of the mouse retina to discover that the majority of synaptic inhibition to the AII amacrine cell (AC), the central neuron in the RB pathway, is provided by a single interneuron type: a multistratified, axon-bearing GABAergic AC, with dendrites in both ON and OFF synaptic layers, but with a pure ON (depolarizing) response to light. We used the nNOS-CreER mouse retina to confirm the identity of this interneuron as the wide-field NOS-1 AC. Our study demonstrates generally that novel neural circuits can be identified from targeted connectomic analyses and specifically that the NOS-1 AC mediates long-range inhibition during night vision and is a major element of the RB pathway.

## Introduction

In dim light, vision originates with rod photoreceptors. In mammals, rod output is conveyed to ganglion cells (GCs), the retinal projection neurons, by the rod bipolar (RB) cell pathway (Demb and Singer, 2015; Famiglietti and Kolb, 1975; Field et al., 2005) (Figure 1). Absorption of photons by rods depolarizes RBs, which therefore are ON cells. RBs make dyad synapses with A17 amacrine cells (ACs) and AII ACs: A17s provide feedback inhibition to the RB [akin to dendro-dendritic inhibition in the olfactory bulb (Isaacson and Strowbridge, 1998; Jahr and Nicoll, 1980, 1982)], and AIIs provide feedforward signals that simultaneously drive excitation and inhibition in ON and OFF GCs, respectively (Murphy and Rieke, 2006; Strettoi et al., 1994; Strettoi et al., 1992).

**Figure 1.**
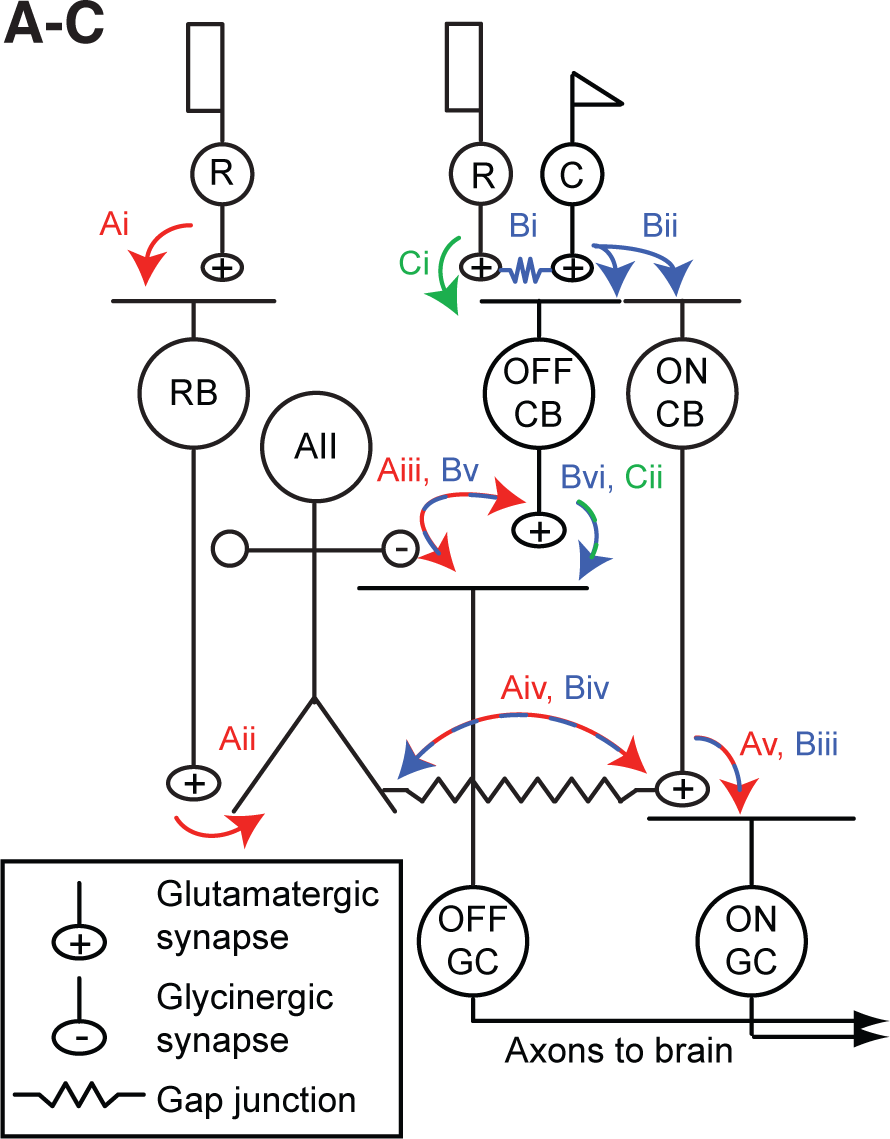
The mammalian retinal pathway for night vision. (Ai-Av) In red: the rod bipolar (RB) pathway of mammalian retina. Rods make synapses onto RBs (Ai), which make synapses onto the AII. (Aii). AIIs make glycinergic synapses (Aiii) onto the terminals of some OFF cone bipolar (CB) cells and onto the dendrites of some OFF ganglion cells (GCs). AIIs are coupled by electrical synapses to the terminals of ON CBs (Aiv), which make glutmatergic synapses onto ON GCs (Av). (Bi-Bvi) In blue: rods are coupled electrically to cones by gap junctions (Bi) and cones make synapses onto ON and OFF CBs (Bii). Depolarization of the ON CB by the cone not only drives glutamatergic transmission to ON GCs (Biii), it also depolarizes AIIs via the electrical synapses (Biv) and thereby elicits glycinergic transmission to OFF GCs and perhaps OFF CBs (Bv). (Ci-Cii) In green: rods make direct chemical synapses onto some types of OFF CB (Ci), which in turn contact OFF GCs (Cii).

Three GC types in the mouse retina receive input from the RB pathway at or near visual threshold: the ON α GC and the OFF α and δ GCs, which exhibit high sensitivity to spatial contrast in the visual scene—similar to primate parasol cells —and project to the geniculo-cortical pathway (Ala-Laurila et al., 2011; Ala-Laurila and Rieke, 2014; Dunn et al., 2006; Grimes et al., 2018a; Grimes et al., 2015; Grimes et al., 2014b; Grimes et al., 2018b; Ke et al., 2014; Kuo et al., 2016; Murphy and Rieke, 2006, 2008). Input from the RB pathway to ON α GCs provides the signal that guides behavior at visual threshold (Smeds et al., 2019).

Signaling from AIIs to ON and OFF pathways is compartmentalized by the morphology of the AII, which is a bistratified cell with distinct neurites in the ON and OFF sublaminae of the inner plexiform layer (IPL; Figure 1). The distal (ON-layer) dendrites receive excitatory input from RBs and make electrical synapses with the axon terminals of ON cone bipolar (CB) cells, particularly type 6 CBs presynaptic to ON α GCs; depolarization of AIIs drives excitatory transmission to ON α GCs (Schwartz et al., 2012). The proximal (OFF-layer) dendrites make inhibitory glycinergic synapses onto the axon terminals of some OFF CBs [primarily type 2 CBs (Graydon et al., 2018)] and some OFF GCs, including OFF α and δ GCs as well as suppressed-by-contrast GCs (Beaudoin et al., 2019; Demb and Singer, 2012; Jacoby et al., 2015). Thus, the AII mediates so-called “cross-over” inhibition, whereby one pathway (ON, in this case) suppresses the other (OFF) and thereby decorrelates their outputs (Demb and Singer, 2015).

Examination of the RB pathway has provided significant insight into the transformation of sensory stimuli into neural responses, particularly under conditions when the signal is sparse (Field et al., 2005). Although well-studied, several uncertainties about RB pathway function remain. Most significantly, we do not know the identity of the spiking, GABAergic AC that drives a receptive field surround in the AII during rod-mediated vision via synaptic inhibition of the AII itself (Bloomfield and Xin, 2000); understanding mechanisms contributing to surround inhibition is important because such inhibition tunes AII responses to spatial features of the visual stimulus.

Here, we identified the major inhibitory input to the AII by combining anatomical, genetic, and electrophysiological analyses in a three-step process. One, we reconstructed ACs that provided synaptic input to AIIs in a volume of mouse retina imaged by scanning block-face electron microscopy [SBEM; (Denk and Horstmann, 2004)]. Two, we evaluated published descriptions of reporter mouse lines to identify genetically-accessible ACs that had the anatomical characteristics of the cells reconstructed from SBEM images. And three, we used electrophysiological recordings and genetics-based circuit analysis to demonstrate that a candidate AC, which provided the great majority of the inhibitory synaptic input to AIIs, exhibited a light response predicted by its anatomy and made GABAergic synapses onto AIIs. This spiking, GABAergic AC is a multistratified, ON AC denoted NOS-1 AC and identified in the nNOS-CreER mouse (Zhu et al., 2014). We conclude that the NOS-1 AC is an integral component of the RB pathway and a significant source of long-range inhibition during night vision. More generally, our study demonstrates the utility of targeted, small-scale “connectomic” analysis for identification of novel neural circuits.

## Results

### AIIs in the mouse retina exhibit a TTX-sensitive, GABAergic receptive field surround

The inhibitory receptive field surround of AIIs has been studied extensively in the rabbit retina, where it is GABAergic and appears to be generated by spiking ACs because it is suppressed by the voltage-gated sodium channel blocker tetrodotoxin (TTX) (Bloomfield and Xin, 2000; Xin and Bloomfield, 1999). Here, we began by probing for the existence of a similar TTX-sensitive surround mechanism in AIIs of the mouse retina (Figure 2A).

**Figure 2.**
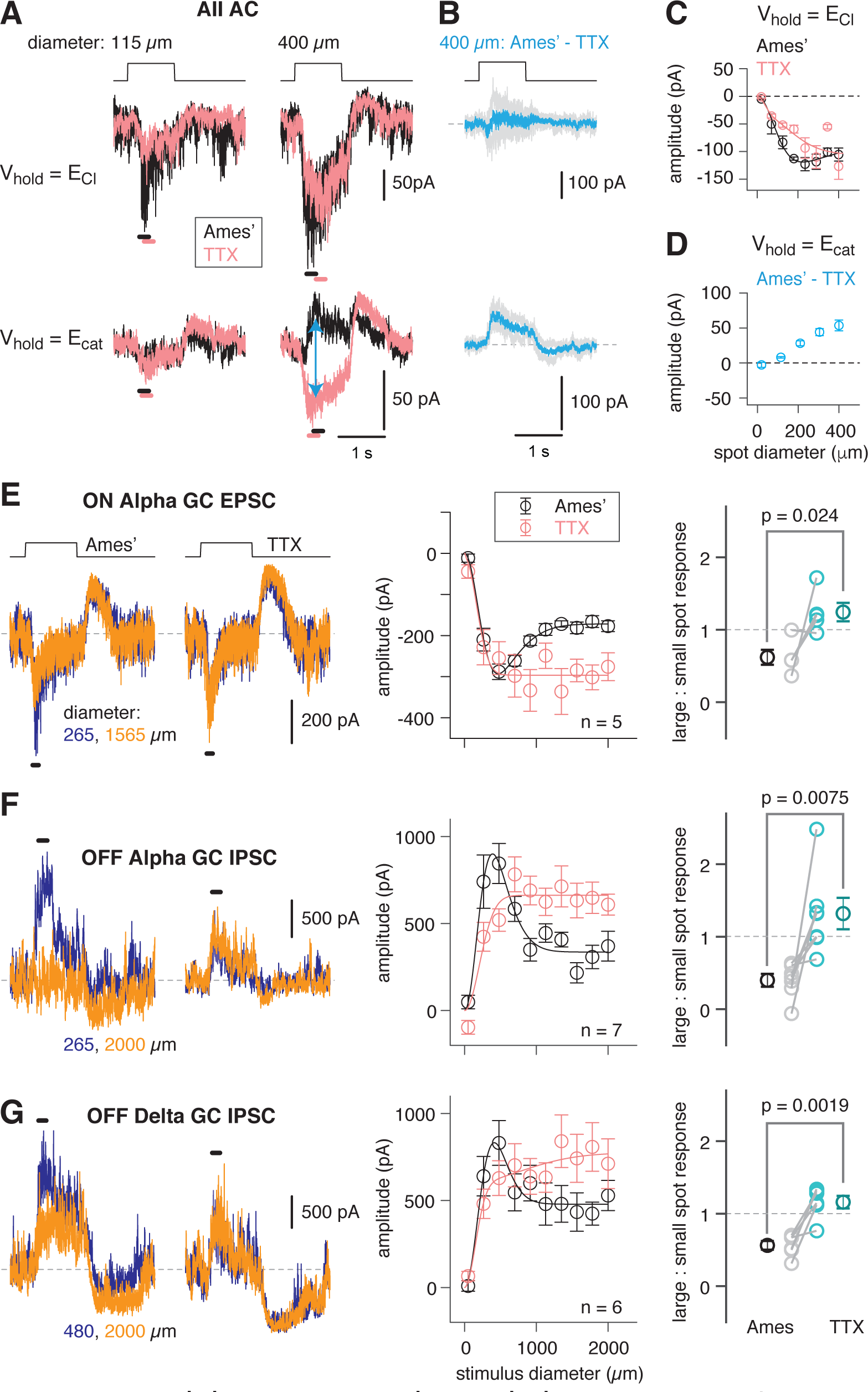
An inhibitory surround recorded in mouse AII ACs. (A) AII responses to spots of light (10 R*/rod/s, 1 s duration) with the indicated diameter in control (Ames’ medium) and after applying TTX (1 µM). Responses were recorded at holding potentials (V_hold_) near E_Cl_ (−70 mV; top row) and near E_cat_ (+5 mV; bottom row). (B) Difference current at each V_hold_ for the 400-µm spot (average of n = 5 cells; shaded areas are ±SEM across cells as a function of time). (C) Response amplitude [measured over a 200 ms window: horizontal bars in (A)] to spot of variable diameter at V_hold_ = E_Cl_ (n = 5 cells). Error bars are ±SEM across cells. (D) Amplitude of difference current at V_hold_ = E_cat_ averaged across cells. Conventions as in (C). (E) Left, light flash (dark background, two spot sizes, 4 or 40 R* / rod / s across cells)-evoked EPSCs in an ON α RGC. Under control conditions, the response is smaller for the larger diameter, illustrating the surround effect, which is blocked by TTX (1 µM). Middle, Spot stimuli of varying diameter (Ames’ and TTX) elicit EPSCs; peak response amplitude measured in a time window (100 – 200 ms; horizontal line at left). Right, ratio of response to large (averaged over the three largest diameters) and small (chosen as the optimal spot size for each cell in the control condition) spots. A ratio < 1 indicates a surround effect. The ratio increased significantly in TTX. (F) Same as (E) for IPSCs in OFF α RGCs (spot intensity, 4 R* / rod /s). (G) Same as (F) for OFF δ RGCs.

AIIs were targeted for recording in the whole-mount retina [see Methods; (Mortensen et al., 2016)], and rods were stimulated with light spots of varying diameter eliciting 10 R* / rod / s. Evoked excitatory currents (voltage-clamp; V_hold_ = E_Cl_ = −70 mV) increased with spot diameter well beyond the physical ∼30 µm width of the AII dendritic field (Figure 2B). This wide receptive field of excitation is explained by electrical coupling within the AII network (Hartveit and Veruki, 2012; Xin and Bloomfield, 1999): an excitatory current originating in surrounding AIIs spreads laterally as a coupling current. Recording at V_hold_ = E_cation_ = +5 mV during spot presentation in control (Ames’) medium yielded an evoked current comprising a mixture of genuine inhibitory current and unclamped coupling current. In the presence of TTX, the inhibitory input was suppressed, leaving the coupling current (Figure 2A). The difference current (Ames’ – TTX; Figure 2A) is the isolated, TTX-sensitive, rod-driven inhibitory input, which increased as a function of spot diameter (Figure 2C). We observed as well that TTX exerted a mild suppressive effect on the excitatory light-evoked current, suggesting that spiking interneurons influence synaptic transmission from the presynaptic bipolar cells (Figure 2B).

### Propagation of the AII surround to downstream ganglion cells

The surround suppression of AIIs is expected to be propagated to the GCs that receive input from the RB-AII network. While recording from three GC types—ON α, OFF α, and OFF δ GCs—rods were stimulated with dim (evoking either 4 or 40 R* / rod / s) spots of varying size; this stimulus will evoke excitatory postsynaptic currents (EPSCs) in ON α GCs and inhibitory postsynaptic currents (IPSCs) in OFF α and δ GCs that reflect the output of the RB-AII network (Murphy and Rieke, 2006, 2008).

As spot diameter increased up to 2000 µm, EPSCs in ON α GCs and IPSCs in OFF α and δ GCs first increased and then decreased in amplitude, reflecting an initial increase in the receptive field center response and then subsequent surround suppression of the center response (Figure 2D-F). In all three cell types, surround suppression was blocked similarly by TTX, suggesting that it was mediated by a common presynaptic mechanism: inhibition of the AII. Thus, the mouse AII exhibits a TTX-sensitive receptive field surround mediated by direct inhibitory synapses and this surround is propagated to GCs. To understand the mechanism for surround inhibition, we searched for the inhibitory ACs presynaptic to the mouse AII.

### Anatomical identification of inhibitory synaptic inputs to AIIs

We began by skeletonizing three AII ACs distributed across serial block face electron microscopy (SBEM) volume k0725 [mouse retina; 50 × 210 × 260 µm; (Ding et al., 2016). AIIs were traced from their locations presynaptic to dyad ribbon synapses at RB terminals that were themselves identified by morphology and position within the inner plexiform layer (IPL) (Graydon et al., 2018; Mehta et al., 2014; Pallotto et al., 2015). For each of these three AIIs, we annotated all of the inputs from ribbon synapses, arising from RBs, as well as all of the conventional synaptic inputs, presumed to arise from inhibitory ACs (Figure 3A). AIIs received substantial input to their distal (ON layer) dendrites from RBs (Table 1; Figure 3B1), consistent with published descriptions (Strettoi et al., 1992; Tsukamoto and Omi, 2013). With regard to inhibitory input, it is notable that virtually all AC inputs to the AII were in the inner IPL (ON layer), on the distal dendrites; the remainder of the AC inputs were on the somas and most proximal portions of the AII dendrites (Table 1; Figure 3B2).

**Figure 3.**
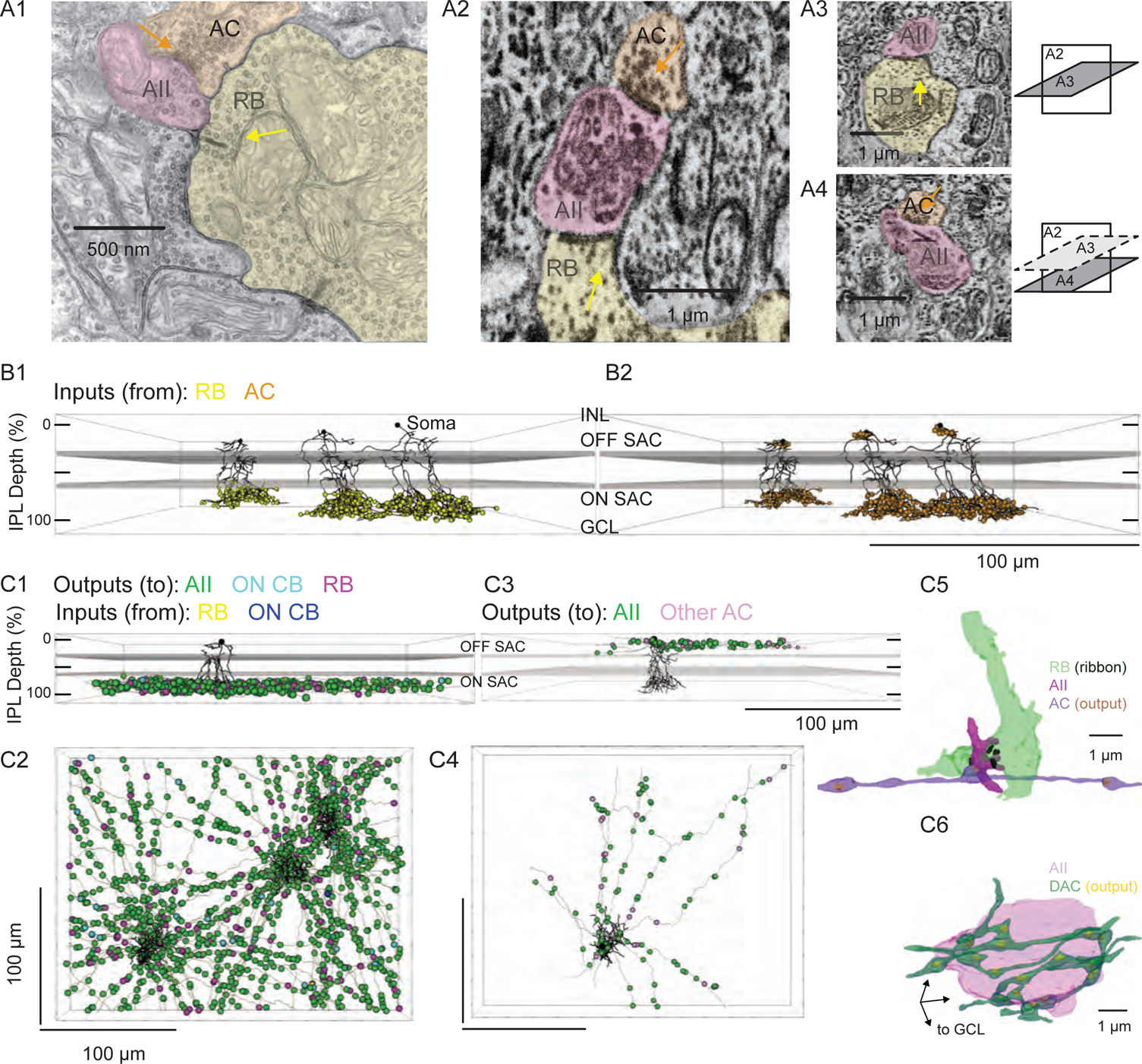
Anatomical characterization of AC axons presynaptic to the AII. (A1) A RB dyad synapse, visualized by TEM. Note the synaptic ribbon (yellow arrow) and the nearby AC input to the postsynaptic AII (orange arrow). (A2) A similar dyad synapse observed by SBEM. Again, the RB ribbon and AC input to the postsynaptic AII are marked with yellow and orange arrows, respectively. (A3, A4) Orthogonal views of the synapse illustrated in A2 showing the RB dyad (top) and the AC input to the postsynaptic AII (bottom). A3 and A4 are from different sections to illustrate the appearance of both ribbon and AC inputs in the section orthogonal to A2. This is illustrated schematicaly by the squares at right. (B) Three AIIs, skeletonized and annotated with either RB ribbon synapse locations (B1) or AC synapse locations (B2). The view is from the side of the volume, representing a transverse section through the retina. The planes containing ON and OFF SAC dendrites are represented by gray rectangles. Note that the majority of AC inputs to the AII are found in the same IPL sublaminae as the RB inputs, the vitreal side of the ON SAC dendrites. As well, AIIs receive no synaptic input from ACs in the IPL sublaminae between the ON and OFF SAC dendrites. (C1) Side view of a single AII and 21 reconstructed presynaptic neurites with output synapses annotated. (C2) An en face view, visualized from the GCL, of 3 AIIs (black) and 61 presynaptic axons. Note the preponderance of green circles, indicating synapses to the three reconstructed AIIs and other AIIs within the volume. Virtually every synapse that was not made with an AII was onto a RB. We observed two synapses onto these axons. (C3) Side view of a single AII and neurites presynaptic to its soma and proximal dendrites. (C4) An *en face* (from GCL) view of the same AII and neurites. (C5) Segmentation of the RB-AII-AC complex. The AC axon is thin, with occasional large varicosities containing clusters of vesicles. It makes synapses with AIIs, usually quite close to RB◊AII ribbon synapses. In this example, the AII dendritic segment receives 4 ribbon inputs from the axon terminal varicosity of a RB and 1 conventional synaptic input from an AC. The other synapses made by this AC axon segment also are with AIIs. (C6) Segmentation of an AII soma and presynaptic neurites, with presynaptic active zones annotated. The image is a tilted side view; the orientation axis (lower left) indicates the relative positions of the IPL and GCL.

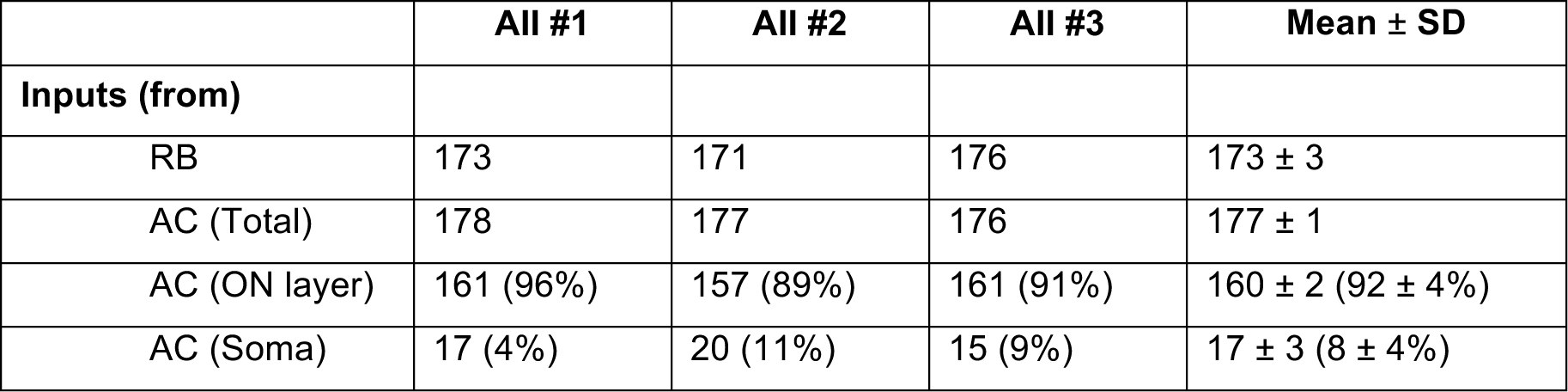

| | All #1 | All #2 | All #3 | Mean ± SD |
| --- | --- | --- | --- | --- |
| **Inputs (from)** |  |  |  |  |
| RB | 173 | 171 | 176 | 173 ± 3 |
| AC (Total) | 178 | 177 | 176 | 177 ± 1 |
| AC (ON layer) | 161 (96%) | 157 (89%) | 161 (91%) | 160 ± 2 (92 ± 4%) |
| AC (Soma) | 17 (4%) | 20 (11%) | 15 (9%) | 17 ± 3 (8 ± 4%) |

For each AII, we skeletonized 21 of the AC inputs to the distal dendrites to assess the morphology of the presynaptic neurons (Figure 3C1,2). Of the 63 AC skeletons created, 61 were of neurites, generally unbranched, that extended through the volume and appeared to be axons: each of these originated from an AC not contained in the SBEM volume (Figure 3C2). After annotating their output synapses, we determined that these axons made synapses with AIIs almost exclusively; the remainder of the output was to RBs with very few synapses to ON CBs and unidentified cells (Table 2; Figure 3C2). This determination was made by tracing the postsynaptic neurites sufficiently to identify RBs from their characteristic axon terminals, which are large and make dyad synapses with presumed AIIs and A17 ACs, and to identify AIIs based on several characteristic features: a soma position at the border of the INL and IPL; very thick proximal dendrites; and a postsynaptic position at RB dyad synapses [see (Graydon et al., 2018; Mehta et al., 2014; Strettoi et al., 1990; Strettoi et al., 1992)].

These 61 axons made 130 synapses onto the 3 reconstructed AIIs, giving rise to ∼25% of the total inhibitory input to each AII. Therefore, assuming that each axon arises from a distinct cell, most AIIs receive ∼2 inputs (i.e. ∼130 synapses/61 axons) from each presynaptic AC; these inputs occur at sites close to RBàAII synapses (Figure 3B, 3C5). The inputs to the AII somas and proximal (OFF layer) dendrites were not considered in as much detail because these were few in number and presumed to arise from dopaminergic ACs (DACs) (Contini and Raviola, 2003; Gustincich et al., 1997; Voigt and Wassle, 1987). Analysis of inputs to one of the three AIIs supported this proposition (Figures 3C3,4): 8 neurites that contacted primarily its soma as well as somas of neighboring AIIs. These 8 neurites made 113 output synapses in total, 90 (80%) of which were to AIIs and 23 were to other cells. AII somas appeared to be enveloped in a “basket” of such neurites (Figure 3C6), as noted previously for DACàAII synapses (Gustincich et al., 1997; Voigt and Wassle, 1987).

Two of the 63 AC skeletons that made inputs to the AII distal dendrites could be traced to a relatively complete neuron contained within the SBEM volume (Figure 4A, B). These two ACs appeared to be of the same type and were characterized as displaced (with somas in the GC layer), multi-stratified cells, with processes in both the ON and OFF strata of the IPL. As well, it was notable that the neurites tended to branch at approximately right angles when viewed *en face*.

**Figure 4.**
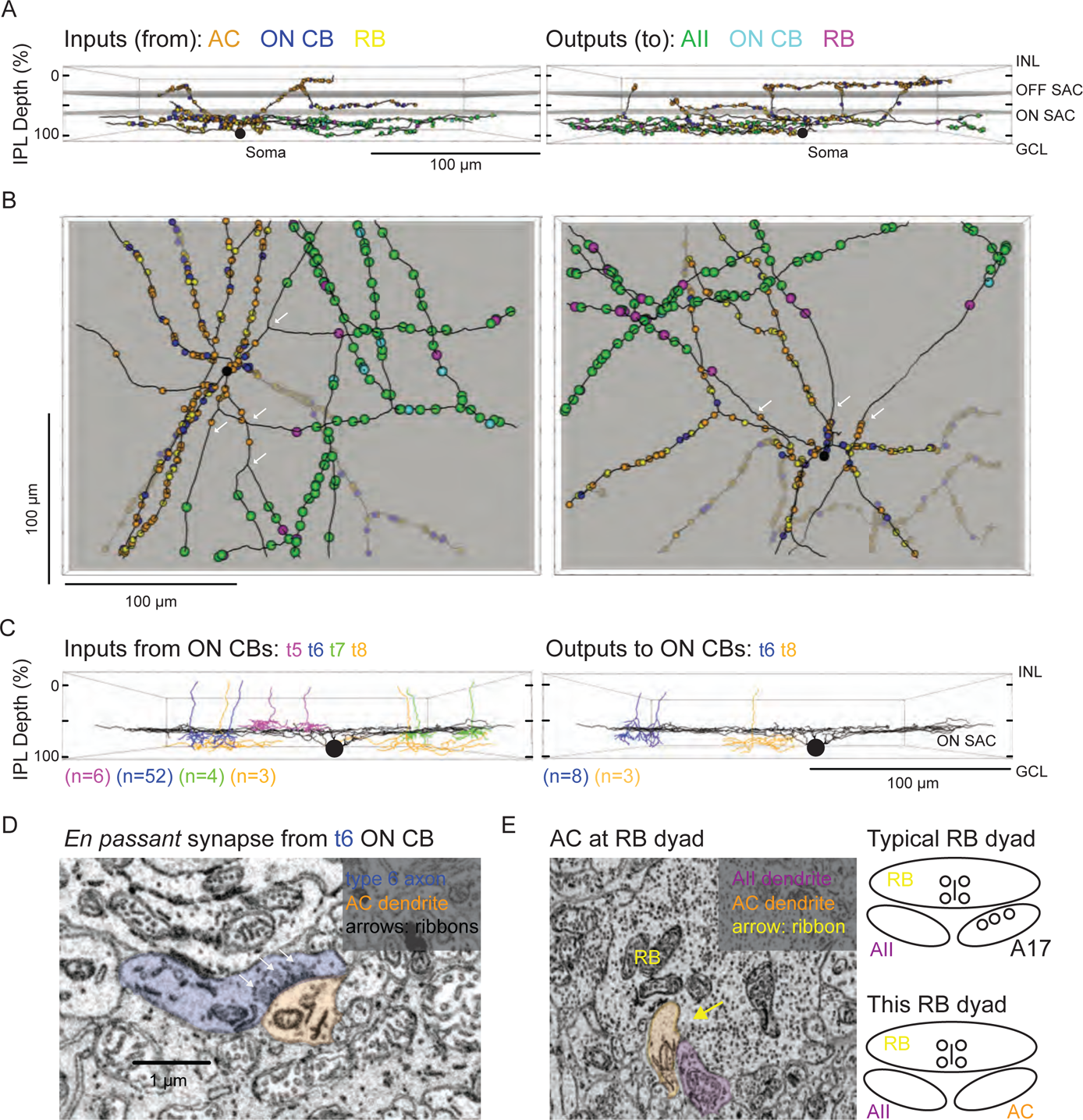
Anatomical characterization of an AC presynaptic to the AII. (A) Skeletonization and annotation of two ACs (left and right panels) with similar morphologies and patterns of synaptic connectivity. Note that both cells, viewed from the side, as in a transverse section through the retina, have similar neurite branching patterns and receive synaptic input from ACs and from ON CBs on dendrites in the OFF laminae of the IPL, on the outer side of the OFF SAC dendrites (i.e., close to 0% IPL depth; here, SAC dendrites are represented as gray rectangles). (B) An en face view (viewed from the GCL; the gray represents the layer of ON SAC dendrites) of the two ACs illustrated in A. Note that their synaptic inputs and outputs are segregated to different sections of their processes; the area receiving input is dendritic, and the area making output is axonal. White arrows indicate areas where dendrites become axons (inputs are proximal to the arrow, closest to the soma; outputs are distal to the arrow, farther from the soma). (C) Side (transverse) view of the retina illustrating an ON SAC (from Ding et al. 2016) and representative ON CBs pre-(at left) or postsynaptic (at right) to the two ACs illustrated in (A) and (B). ON CBs were classified based on axon branching pattern and stratification depth relative to the ON SAC dendrites. (D) Example en passant ribbon-type synapses in a type 6 ON CB axon. Note three ribbons clustered together and presynaptic to the same AC process. (E) Example of RB dyad at which the AC type shown in (A) and (B) replaces the A17 as one of the two postsynaptic cells (see schematic at right).

### Anatomical assessment of an AC circuit presynaptic to the AII

We annotated all of the synaptic connections—inputs and outputs—made by the two ACs contained within the volume and then identified each pre- or postsynaptic partner (Figure 4A, B). Both ACs exhibited very similar patterns of connectivity, as quantified in Table 2. Interestingly, the vast majority of synaptic output was to AIIs (confirmed as described above); a smaller but significant portion was to RBs (again, confirmed as above), with the remainder to ON CBs (which were skeletonized completely; below). We did not observe any synapses onto other ACs or onto GCs. Thus, these two ACs appear to be representatives of a single AC type that contacts preferentially neurons in the RB pathway, specifically AIIs. Given the similarity of the postsynaptic target neuron populations of these ACs and the axons reconstructed partially and described above (Figure 3C1, C2), we believe all to be representative of the same AC type, a cell that contacts AIIs preferentially and provides the vast majority of inhibitory synapses to the AII.

Both ACs received very similar numbers of inputs from conventional and ribbon synapses (Table 2). The conventional synapses were presumed to arise from inhibitory ACs; these synapses were annotated, but we did not attempt to identify the presynaptic cells. Excitatory inputs both from *en passant*, axonal ribbon synapses and from more typical, axon terminal ribbon synapses were observed; analysis of the presynaptic cells revealed them to be ON bipolar cells—both RBs and ON CBs—exclusively (Table 2). Thus, this AC appears to receive only excitatory ON bipolar input despite its being a multi-stratified cell with processes in both ON and OFF sublaminae of the IPL.

|  | **Axons**<br>**(Figure 3C2)** | **Cell 1**<br>**(Figure 4)** | **Cell 2**<br>**(Figure 4)** |
| --- | --- | --- | --- |
| **Inputs (from)** | Total: 2 | Total: 314 | Total: 297 |
| AC |  | 200 (63.5%) | 183 (61%) |
| RB | 1 | 47 (15%) | 56 (19%) |
| ON CB | 1 | 66 (21%) | 55 (18.5%) |
| Unidentified |  | 2 (<1%) | 3 (1%) |

| Outputs (to) | Total: 1425 | Total: 115 | Total: 106 |
| --- | --- | --- | --- |
| All | 1212 (85%) | 93 (81%) | 86 (81%) |
| RB | 173 (12%) | 11 (9.5%) | 18 (17%) |
| ON CB | 35 (2%) | 11 (9.5%) | 2 (2%) |
| Unidentified | 5 (<1%) |  |  |

The 121 ON CB synapses onto the 2 ACs arose from 67 presynaptic ON CBs, and we skeletonized these ON CBs in order to identify them based on morphological characteristics and axon terminal depth within the IPL. We used the positions of the cholinergic starburst ACs in the SBEM volume as standard markers of IPL depth (Figure 4C) (Ding et al., 2016; Helmstaedter et al., 2013; Manookin et al., 2008; Sabbah et al., 2017; Stabio et al., 2018). Most of the presynaptic ON CBs belonged to the type 6 population; the others were a mix of types 5, 7 and 8 CBs (two cells could not be fully reconstructed and were unidentified) (Figure 4C). Fifteen *en passant* (axonal) synapses onto the outer (OFF-layer) dendrites of the reconstructed ACs were observed (Figure 4D): 13 from type 6 cells and two from the unidentified cells (likely type 6 CBs). Both the occurrence of these axonal ON bipolar cell synapses and their general appearance are consistent with previous reports (Dumitrescu et al., 2009; Hoshi et al., 2009; Kim et al., 2012; Lauritzen et al., 2013). Although, we observed *en passant* synapses in the axons of some, but not all, reconstructed type 5, 7 and 8 ON CBs, none of these *en passant* synapses were presynaptic to the reconstructed ACs. Additionally, *en passant* synapses in axons of type 6 cells had a markedly distinct appearance: ribbons that tended to occur in clusters of at least three, all apposed to the same postsynaptic process (Figure 4D). In this respect, they resemble strongly the axonal ribbons observed in calbindin-positive ON CBs of the rabbit retina, which are likely homologous to mouse type 6 CBs because both share similar morphology and make synapses with ON α GCs (Hoshi et al., 2009; Kim et al., 2012; Schwartz et al., 2012; Tien et al., 2017).

The 103 RB synapses with these two ACs arose from 77 RBs (all of which were skeletonized; not shown). All RBàAC synapses were dyads (Figure 4E); at 75% of these dyads, the AC replaced the A17 (i.e. the other postsynaptic cell was an AII), and at the remainder of the dyads, the AC either replaced the AII (18%) or else the identity of the second postsynaptic cell could not be confirmed (7%). In at least two of the cases in which the second cell at the dyad could not be identified, the dendrite had the appearance of those of the skeletonized ACs: a very thick dendrite containing clear cytoplasm. Thus, it appears that this AC receives a significant portion of its excitatory input (48%; see Table 2) from the RB population but that any individual RB provides only one or two synapses to a single AC; the latter finding is consistent with the vast majority of RB output being at dyad synapses with AII and A17 amacrine cells (Demb and Singer, 2012).

A number of multistratified AC types have been reported in anatomical studies of the mouse retina (Badea and Nathans, 2004; Lin and Masland, 2006; Perez De Sevilla Muller et al., 2007). The vast majority of these, however, differ in their morphology from the ACs studied here: e.g., of the 16 wide-field, axon-bearing ACs categorized by Lin and Masland (2006; their Figure 10), none have axons in the inner (ON) layer of the IPL and a narrow field of dendrites in the outer (OFF) layer. We were struck, though, by the resemblance of the AC identified here to one identified in a screen of Cre-driver lines: the NOS-1 AC of the nNOS-CreER mouse (Zhu et al., 2014). Therefore, we tested the hypothesis that the NOS-1 AC is the spiking, ON AC that provides inhibitory synaptic input to the AII.

### The NOS-1 AC is a spiking ON cell

NOS-expressing (NOS+) ACs were targeted for *in vitro* recording by crossing nNOS-CreER mice with Cre-dependent reporter mice (Ai32: ChR2/eYFP fusion protein expression; Zhu et al., 2014; Park et al., 2015) and then inducing recombination in ∼1 month old offspring by tamoxifen injection (see Methods). eYFP fluorescence (eYFP+) observed by two-photon laser scanning microscopy (2PLSM) was used to target NOS+ ACs for recording. We studied retinas in which Cre expression was induced robustly; these had 110 ± 5 cells mm^-2^ (mean ± sem) labeled in the GCL and 128 ± 5 cells mm^-2^ labeled in the INL (n = 7 retinas from 7 mice). Based on an earlier description of this driver line, we assumed that labeled cells in the GCL included a mixture of NOS-1 and NOS-2 ACs, with NOS-1 cells having the bistratified morphology described here and NOS-2 cells exhibiting thick, spiny dendrites that project into the central level of the IPL, between the layers marked by dendrites of ON and OFF starburst ACs (Jacoby et al., 2018; Zhu et al., 2014). Labeled cells in the INL included additional NOS-2 ACs (Zhu et al., 2014) as well as other AC types that projected into the outer most levels of the IPL but were not studied here.

Dye filling (Lucifer Yellow) of recorded eYFP+ cells in the GCL revealed that these cells most typically were NOS-1 ACs, with bistratified dendrites and long axons identified by 2PLSM following recording and, in some cases, by subsequent analysis by confocal microscopy (n = 13 cells; Figure 5A). Whole-cell recordings of membrane voltage (i.e. current-clamp recordings) showed that NOS-1 cells fired spikes in response to positive contrast and that spiking could be suppressed completely by negative contrast (Figure 5B). Responses increased in magnitude with increasing spot diameter, suggesting an integration area of at least ∼500-µm diameter (Figure 5C). NOS-2 cell membrane voltage responses to light clearly differed from those of NOS-1 cells (Figure 5D, E). NOS-2 cells were non-spiking with graded, depolarizing responses to both positive and negative contrast and are therefore ON-OFF cells (Jacoby et al., 2018). Both ON and OFF responses of NOS-2 cells increased with spot diameter, with a relatively more gradual increase for the OFF response (Figure 5F).

**Figure 5.**
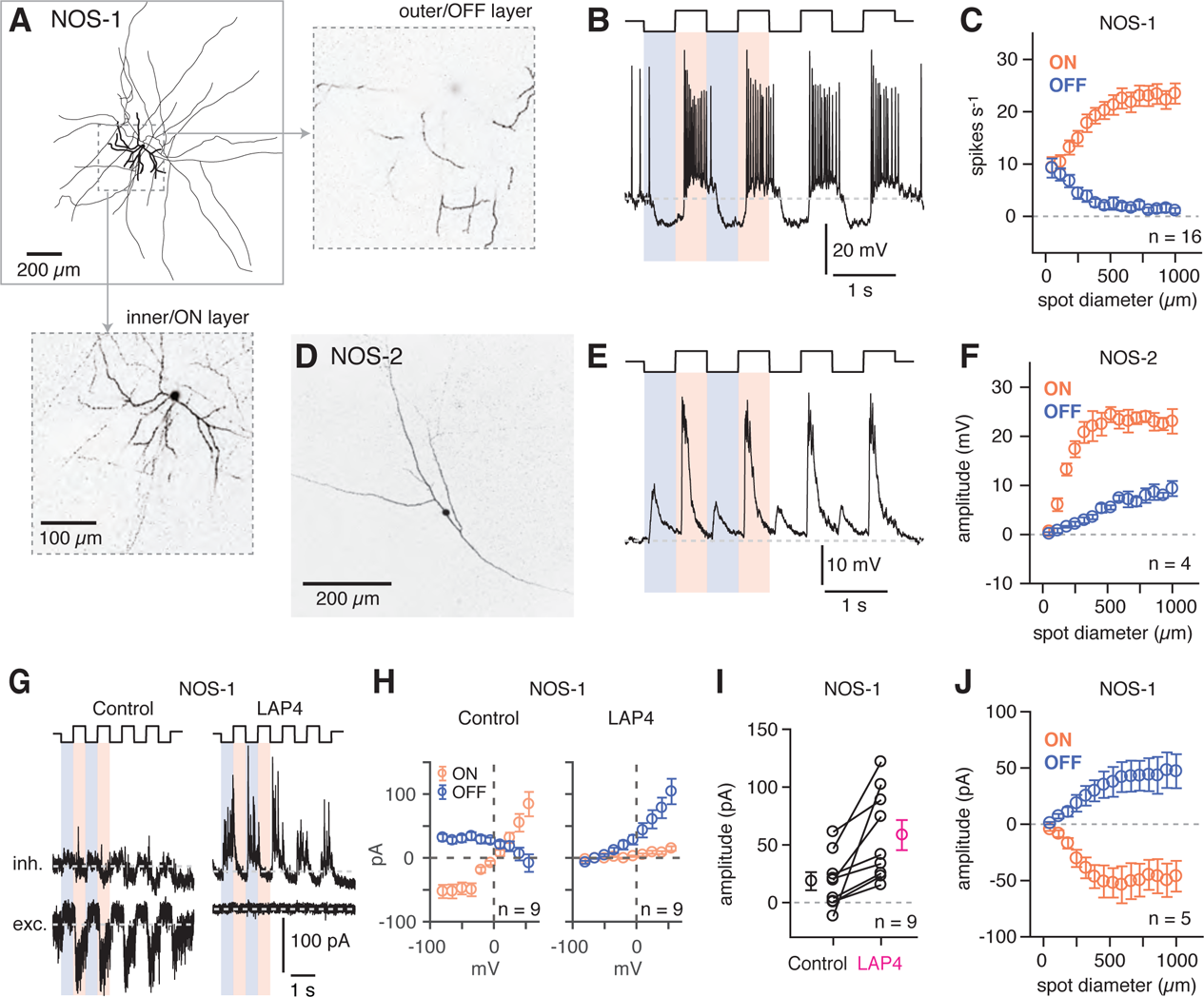
NOS-1 ACs are spiking ON-center cells that can be distinguished from NOS-2 ACs. (A) Morphology of a NOS-1 AC. Dendrites (thick) and axons (thin) were drawn from confocal images of a NOS-1 AC filled by Lucifer Yellow during whole-cell recording. Single confocal sections are shown for the inner/ON and outer/OFF layers of processes for the region indicated (dashed, boxed region). The cell is bistratified in the region proximal to the cell body; only the ON-layer processes are shown in the drawing. (B) Membrane potential recording for the cell in (A). The cell had a baseline firing rate at mean luminance (∼10^4^ R*/cone/s) that modulated above and below baseline during positive and negative contrast periods, respectively (spot diameter, 600 µm; 100% contrast). (C) Population (n = 16 cells) changes in firing rate during positive contrast (ON response) and negative contrast (OFF response) as a function of spot size (100% contrast). Firing rate was computed over a 500-ms time window for each contrast. Error bars indicated ±SEM across cells. (D) Collapsed confocal stack (maximum projection image) of a filled NOS-2 cell. (E) The NOS-2 cell in (D) responds with depolarization at both positive and negative contrast, an ON-OFF response (spot diameter, 600 µm; 100% contrast). (F) Population (n = 4 cells) changes in membrane potential as a function of spot size (measured over a 100-ms time window). Conventions are the same as in (C). (G) Excitatory and inhibitory current measured in a NOS-1 cell to a spot stimulus (diameter, 400 µm). After adding L-AP4 (20 µM) to suppress the ON pathway, the excitatory current is blocked and the inhibitory current increases and is OFF responding. (H) Current-voltage (I-V) plots for ON and OFF responses for data in (G) averaged across cells (measured over a 100- to 200-ms time window). Error bars indicate ±SEM across cells. (I) Population change in the OFF inhibitory current after adding L-AP4. Individual cell data are connected by lines. Population data indicate mean ±SEM across cells. (J) Excitatory current amplitude for a population of NOS-1 cells (n = 5 cells) as a function of spot diameter (100% contrast; measured over a 100-ms time window). Error bars indicate ±SEM across cells.

Voltage-clamp recording from NOS-1 cells demonstrated that positive contrast evoked excitatory synaptic input, measured as inward current relative to a standing current (V_hold_ = E_Cl_) (Figure 5G, H). Negative contrast evoked a net outward current, consistent with temporary suppression of ongoing presynaptic glutamate release (Figure 5G, H, J). Excitatory input was blocked completely by L-AP4, which suppresses ON bipolar cells (Slaughter and Miller, 1981) (Figure 5G). Inhibitory synaptic input (V_hold_ = E_cat_) was measured at both positive and negative contrast but was typically small under control conditions; in the presence of L-AP4, however, inhibition persisted only in response to negative contrast and became larger than in control conditions (Figure 5G, H); the amplitude of the response to negative contrast in L-AP4 increased by 38 ± 11.4 pA (t = 3.3; n = 9; p < 0.005; Figure 5I). Thus, NOS-1 cells receive tonic glutamatergic input from ON bipolar cells that is modulated by contrast and inhibitory inputs from both ON and OFF pathways.

### The NOS-1 AC is the predominant NOS-expressing AC in the ganglion cell layer and is distinct from CRH-expressing ACs

To determine the relative density of displaced NOS-1 and NOS-2 ACs in the GCL of the nNOS-CreER retina, we made loose patch recordings of responses to light from displaced eYFP+ ACs targeted by 2PLSM (Figure 6A). Most displaced ACs were spiking cells with sustained ON responses, confirming their identity as NOS-1 cells (n = 22/26 cells, three retinas from two mice). One exception was a spiking cell with both ON and OFF responses (Figure 6A, asterisk); we did not study this cell further, but it likely represents either a subtype of nNOS AC that is primarily in the INL and only rarely displaced to the GC or a case of ectopic Cre and / or reporter expression. In the NOS-1 population, ∼2-3 cells could typically be found within ∼54,500 µm^2^, the cross-sectional area of our SBEM volume; this cell density is consistent with our finding two putative NOS-1 cells in the SBEM data set.

**Figure 6.**
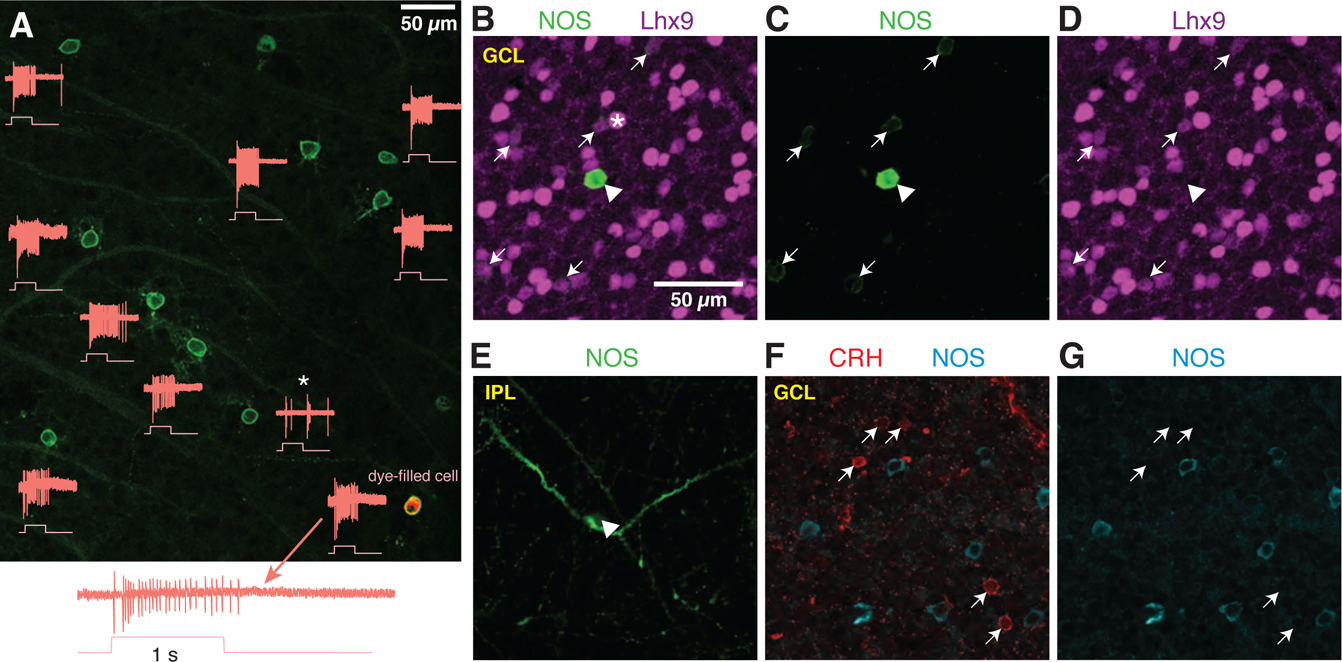
NOS-expressing ACs in the ganglion cell layer are primarily NOS-1 cells. (A) Loose-patch spike recording from a region of retina with cells labeled in the nNOS-CreER x Ai32 mouse. The response to a light flash (800 µm-diameter, ∼10^4^ R*/cone/s) is shown next to each soma that was recorded. The majority of cells showed a sustained ON response in the spike rate during the light flash. In one case, the response differed (*) and showed a transient ON-OFF response. The cell at lower right was subsequently studied by whole-cell recording and was filled with dye; for this cell, the spike response is shown at an expanded scale below the image. (B-E) P12 retinas (C57/B6 wild-type) stained with antibodies to label NOS and Lhx9 expressing cells in the GCL. Most NOS-expressing cells showed dim labeling for Lhx9 antibody (arrows). A well-stained NOS-expressing cell in the center (arrow-head) did not show Lhx9 labeling. Laser power was increased for Lhx9 imaging, such that fluorescence of strongly-labeled cells was saturated (*, example cell), making it easier to visualize weakly-labeled cells. NOS labeling alone (C) and Lhx9 labeling alone (D). (E) Same as (C) with the image plane shifted to the inner plexiform layer (IPL). The NOS-expressing cell that lacked Lhx9 expression (arrowhead) had thick dendrites that could be followed into the IPL, with the characteristic properties and stratification of a NOS-2 cell. (F-G) P12 retina (C57/B6) stained with antibodies to label corticotropin releasing hormone (CRH) and NOS. NOS-expressing cells do not overlap with CRH-expressing cells. NOS labeling alone (G). Scale bar in (B) applies to (B) – (G).

Additionally, we examined NOS-expressing cells by nNOS antibody labeling during a developmental period (P8 – P14) during which a subset of NOS+ cells express the transcription factor Lhx9 (Balasubramanian et al., 2017). Two populations of NOS+ cells were observed: a small group of brightly-labeled cells (n = 19 cells) and a much larger group of dimly-labeled cells (n = 550 cells; six retinas from six mice; n = 2 retinas each at P8, P12 and P14). In retinas double-labeled with the Lhx9 antibody (two retinas from two animals, P12), only the dimly-labeled NOS+ cells were Lhx9+ (Figure 6B-D) (n = 146/156 cells). The brightly-labeled, Lhx9-cells (n = 5/5 cells, two retinas from two mice) exhibited dendrites that could be followed into the center of the IPL, where they gave rise to the thick, spiny processes characteristic of NOS-2 cells (Figure 6E). We conclude that Lhx9 expression distinguishes NOS-1 from NOS-2 cells in the GCL at P12, and that at all developmental time points, NOS-2 cells are rarely found in the GCL (3.3% of cells), making NOS-1 cells a large majority in the GCL (96.7% of NOS+ cells).

The NOS-1 cell was described also as the CRH-2 cell (Corticotropin-releasing hormone AC-2) based on labeling in the CRH-ires-Cre retina (Zhu et al., 2014). We, however, found that Cre-expressing cells in the CRH-ires-Cre::Ai32 retina are rarely labeled by the nNOS antibody (Park et al., 2018), suggesting limited overlap in CRH+ and NOS+ AC populations. Indeed, we found no co-expression of CRH and NOS by immunohistochemistry (P12 retina) (n = 66 CRH+ cells and 85 NOS+ cells, one retina) in neurons in the GCL (Figure 6E-G). Thus, NOS-1 ACs do not appear to co-express CRH, and we conclude, then, that Cre expression in NOS-1 ACs of the CRH-ires-Cre line is rare and unsystematic.

### Confirmation of synapses between NOS-1 cells and AIIs

To demonstrate that NOS-1 ACs provide synaptic input to AIIs, we eliminated Cre-expressing ACs in the nNOS-CreER retina by intraocular injection of an AAV containing a Cre dependent-DTA construct followed 2-3 days later by tamoxifen administration. Four weeks later, we stimulated the retina *in vitro* with dim (40 R* / rod / s) spots of varying size and recorded light-evoked currents in AIIs at V_hold_ = E_Cl_ or V_hold_ = E_cat_ (Figure 7A; as in Figure 2A). Following ablation of NOS+ ACs (including both NOS-1 and NOS-2 cells), the recorded excitatory currents were smaller than in the control condition and were unaffected by TTX (Figure 7B; compare to Figure 2B), consistent with a loss of inhibitory input from NOS-1 ACs to presynaptic RBs (Figure 4 and Table 2). Most significantly, though, we observed a lack of inhibitory TTX-sensitive surround in AII responses following NOS+ AC ablation (Figure 7A, C): the TTX-sensitive IPSC evoked by a 400-µm diameter spot was significantly smaller in DTA-expressing retinas than in controls (compare to Figure 2).

**Figure 7.**
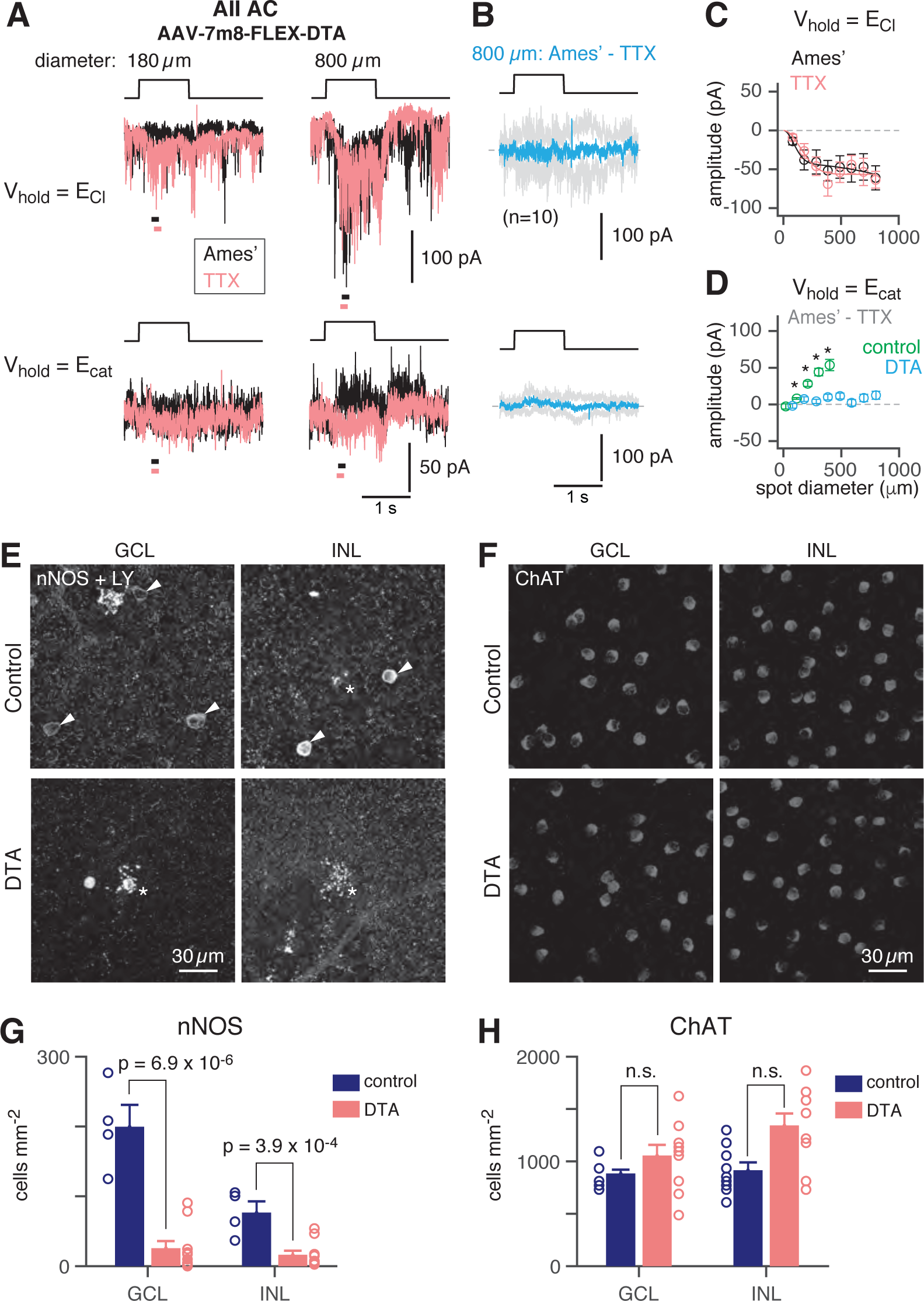
NOS+ ACs generate the TTX-sensitive receptive field surround of the AII. (A) AII responses to spots (diameter indicated; 10 R*/rod/s, 1 s duration) in control (Ames’) and in TTX (1 µM) recorded at E_Cl_ (−70 mV; top) and near E_cat_ (+5 mV; bottom) in a nNOS-CreER retina injected with Cre-dependent DTA virus (AAV-7m8-FLEX-DTA). (B) Difference current at each V_hold_ for the 800-µm diameter spot (average of n = 5 cells; shaded areas are ±SEM across cells as a function of time). (C) Spot (variable diameter) response amplitudes [measured over a 200-ms time window, indicated by horizontal bars in (A] at V_hold_ = E_Cl_ (n = 5 cells). Error bars are ±SEM across cells. (D) Difference current amplitude at V_hold_ = E_cat_ averaged across cells. Same conventions as in (C). Recordings from control retinas (no DTA virus; from Figure 2) superimposed. Responses to similar diameter spots were significantly smaller in the DTA group compared to control (one-tailed t-tests, *): 80/115-µm, t = 2.12, p = 0.027; 180/210-µm, t = 3.11, p = 0.0041; 285/305-µm, t = 2.5, p = 0.0083; 385/400-µm, t = 2.56, p = 0.012. (D) nNOS immunolabeling in GCL and INL, centered on a region with a recorded AII (visible in some images, marked by *). (E) Same format as (D) for ChAT immunolabeling of starburst ACs. (F) NOS+ cell density over a square region (0.64 x 0.64 mm) centered on a recorded AII and visualized by nNOS immunolabeling: DTA vs. control (Cre-positive with no DTA virus, n = 3; or Cre-negative with DTA virus, n = 1). Virus-injected retinas had significantly fewer cells (one-tailed t-test): GCL, t = 7.03, p = 6.9 x 10^-6^; INL, t = 4.46, p = 3.9 x 10^-4^. (G) Same format as (F) for ChAT immunolabeling. Cell density assessed over a square region (0.16 x 0.16 mm) centered on a recorded AII. Starburst AC density in DTA virus-injected retinas was no smaller than in controls.

We confirmed the elimination of NOS+ ACs by nNOS antibody staining and found that the number of NOS+ cells was reduced strongly in DTA-expressing retinas relative to controls (Figure 7D, F). To confirm that DTA expression did not result in non-specific ablation of ACs, we used ChAT antibody immunolabeling to quantify the number of starburst ACs (in both the INL and GCL) in control and DTA-expressing retinas and found no reduction in the experimental group (Figure 7E, G). Thus, elimination of NOS+ ACs provides evidence that a NOS+ cell, most likely the NOS-1 AC, is necessary for generating the AII inhibitory surround.

To confirm that NOS-1 ACs provide synaptic input to AIIs, we performed optogenetic circuit-mapping experiments after inducing ChR2/eYFP expression in NOS+ cells, as described above. After blocking the influence of the photoreceptors (see Methods), we recorded from cells in a retinal whole-mount preparation and stimulated ChR2-expressing neurons with bright blue light. A representative ChR2-expressing NOS-1 cell (n = 3 total) responded within a few ms of the optogenetic stimulus with increased spiking (Figure 8A); IPSCs recorded in AIIs (normalized to their peak amplitude, 39 ± 6 pA) were observed a few ms later (Figure 8A), consistent with a monosynaptic connection from NOS-1 cells.

**Figure 8.**
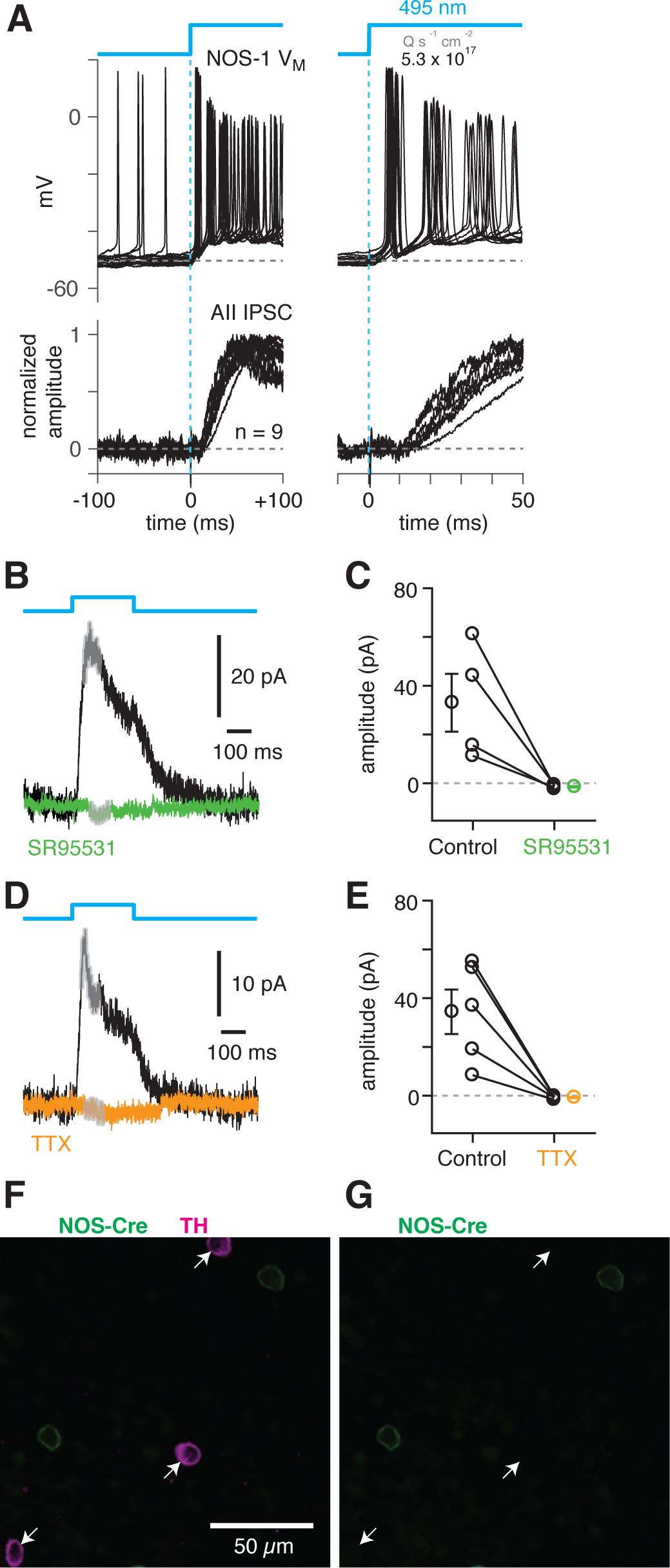
NOS-1 cells make synapses with AII amacrine cells. (A) Top/Left, optogenetic stimulation of a ChR2-expressing NOS-1 cell (of n = 3 total) in the nNOS-CreER::Ai32 retina responded with increased spike firing to blue light (n = 12 trials overlaid). Response was recorded in whole-mount retina in the presence of drugs to block photoreceptor-mediated inputs to retinal circuitry: DNQX (50 µM), D-AP5 (50 µM), L-AP4 (2 µM), and ACET (1 µM). Bottom/Left, the optogenetic stimulus evoked IPSCs (V_hold_ = E_cat_;) in AII ACs (n = 9 cells). Responses are normalized to the maximum amplitude, 39 ± 6 pA (measured over a 60-70 ms time window). Right, expanded version of traces at left. The initial spike in the NOS-1 cell occurred a few milliseconds after optogenetic stimulation top), followed a few milliseconds later by the onset of the AII IPSCs. (B) In AII ACs recorded under the conditions in (A), inhibitory current (measured over the gray region) was blocked by the GABA-A receptor antagonist SR95531 (50 µM). (C) Effect of SR95531 in a sample of AII ACs (n = 4 cells). Error bars indicate ±SEM across cells. (D) Same format as (B) with the sodium channel blocker TTX (1 µM). (E) Same format as (C) with TTX (n = 5 cells). (F) Confocal image of the inner nuclear layer of a retina from the nNOS-creER:: Ai32 mouse. A tyrosine-hydroxylase (TH) antibody was used to label dopaminergic ACs (arrows), which did not overlap with Cre-expressing NOS+ ACs. (G) Same image as (F) without the TH labeling. None of the cells with TH immunolabeling (arrows) were eYFP+.

ChR2-evoked IPSCs in AIIs were blocked by SR95531 (50 µM) and therefore were mediated by GABA_A_ receptors (Figure 8B, C) (reduction of 34.5 ± 11.7 pA, or 108 ± 4%; t = 28.5; n = 4; p < 0.001). In a second group of cells, the IPSCs were blocked by TTX (1 µM), indicating that they arose from a spiking presynaptic AC (Figure 8D, E) (reduction of 35.1 ± 8.9 pA or 105 ± 4%; t = 29.5; n = 5; p < 0.001). These results are consistent with a direct synaptic input from the NOS-1 AC to the AII because input from the non-spiking NOS-2 AC would not be TTX-sensitive. The spiking dopaminergic AC (DAC) also makes GABAergic synapses with the somas and proximal dendrites of AIIs (Figure 3B2, C3-5) (Gustincich et al., 1997), but the DACs in the nNOS-CreER retina do not express ChR2/eYFP, as demonstrated immunohistochemically: there was no overlap between TH+ cells, identified by TH immunolabeling (n = 90 cells), and eYFP+ cells (n = 198 cells, five retinas from three mice) (Figure 8F, G).

Additionally, we recorded ChR2 evoked IPSCs in RBs in retinal slices (n=2) to confirm the functionality of NOS-1 ACàRB synapses observed in our anatomical analyses (Figures 3C1, C2, C6 and 4A, B). Here, we had to include the K channel blockers 4-AP and TEA in the external solution to enhance the excitability of cut NOS-1 AC axons so that they could be stimulated adequately in the slice (Figure 9, left); control experiments recording from AIIs in both slice (Figure 9, left) and whole-mount (Figure 9, right) retinal preparations demonstrated that this manipulation enhanced release from NOS-1 cells significantly (n=3). Notably, enhancing excitability in both retinal slices and whole-mount preparations did not change the latency of the IPSCs, supporting the conclusion that they are monosynaptic.

**Figure 9.**
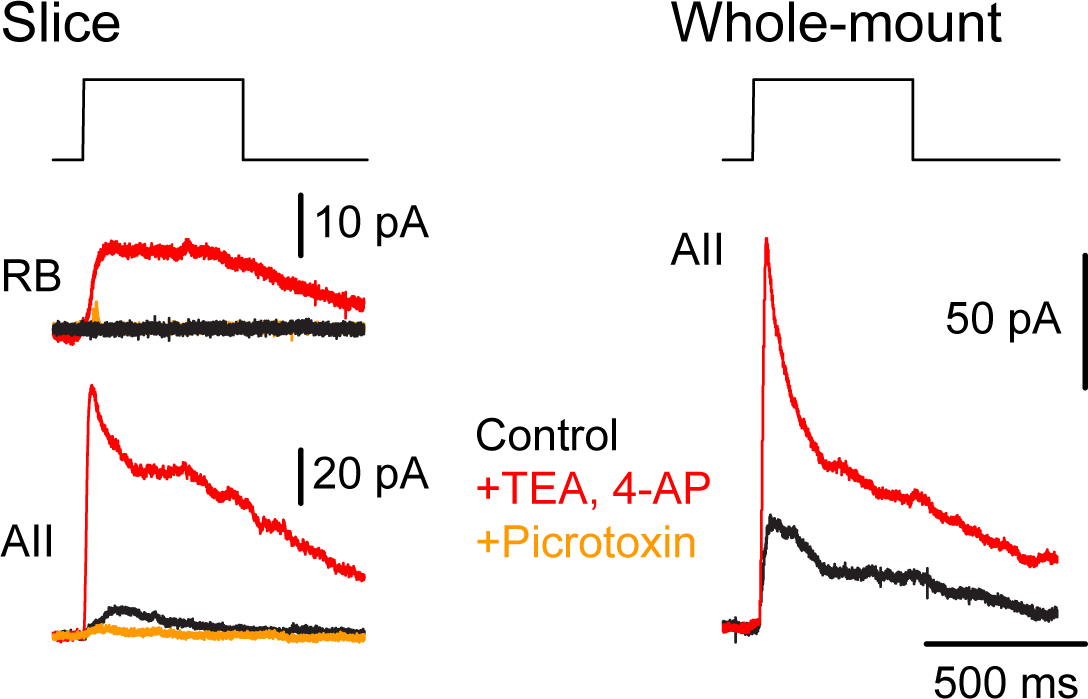
NOS-1 ACs make synapses with RBs. At left, recordings from a RB (top) and an AII (bottom) in a retinal slice demonstrate that optogenetic stimulation of cre-expressing cells in the nNOS-CreER::Ai32 retina evoked inhibitory currents (V_hold_ = E_cat_). Potentiation of presynaptic depolarization with K channel blockers was necessary to elicit responses in RBs owing to the small number of presynaptic axons preserved in the 200-µm thin slice. At right, K channel blockers potentiate larger inhibitory currents (V_hold_ = E_cat_) recorded in an AII and evoked by optogenetic stimulation of cre-expressing cells in a whole-mount preparation of nNOS-CreER::Ai32 retina.

## Discussion

Ultrastructural analysis of inhibitory input to three AIIs indicated that the majority of AC synapses arise from a single type of displaced, multistratified, wide-field cell that contacts AIIs preferentially. Two relatively complete examples of this AC were found within our SBEM volume, and examination of this cell type’s outputs revealed that ∼80% of its synapses were presynaptic to AIIs, with the majority of the remainder presynaptic to RBs. This AC received excitatory input exclusively from ON bipolar cells, both at en-passant axonal synapses in the OFF strata of the IPL and at axon terminal synapses in the ON strata of the IPL. The AC identified by SBEM analysis is a morphological match to the genetically-identified NOS-1 AC, which we demonstrated to have physiological functions predicted by the ultrastructural analysis: it is an ON AC that provides GABAergic inhibition to AIIs and RBs. We therefore consider the NOS-1 AC to be an integral part of the well-studied mammalian RB pathway, serving as the major source of direct, long-range inhibition during night vision.

### Multiple cell types in nNOS-creER retina

Cre-mediated recombination drove ChR2/eYFP reporter expression in multiple AC types in the nNOS-creER retina. We observed two AC types, called NOS-1 and NOS-2, (Jacoby et al., 2018; Zhu et al., 2014), consistent with reports of nNOS expression in at least two AC types (Chun et al., 1999; Kim et al., 1999).

The NOS-1 AC is described above. The NOS-2 AC is a monostratified cell, with dendrites in the center of the IPL (between the processes of ON and OFF starburst ACs; Figure 5D); its soma is either in the conventional location in the INL (∼75%) or displaced to the GCL (∼25%) (Chun et al., 1999; Jacoby et al., 2018; Kim et al., 1999; Zhu et al., 2014). There were additional labeled ACs in the INL of the nNOS-CreER retina: their processes stratified in the OFF layers proximal to the INL, but the cells were not characterized further.

Our anatomical evidence, however, supports the conclusion that the NOS-1 AC is the only Cre-expressing AC presynaptic to the AII: NOS-2 dendrites are confined to the center of the IPL, between the OFF and ON starburst ACs, whereas inhibitory inputs to AIIs are either distal to the OFF starburst processes and apparently arise from dopaminergic ACs (not labeled in the nNOS-CreER line; Figure 6F, G) or proximal to the ON starburst AC processes (Figure 3). As well, in tracing many AC inputs to AIIs, composing ∼25% of the total inhibitory synaptic input to these AIIs, we never encountered an axon that stratified between the ON and OFF starburst ACs.

### The NOS-1 AC is the dominant inhibitory input to the AII and generates the AII surround

The AII has an ON-center receptive field with an antagonistic surround (Bloomfield and Xin, 2000; Nelson, 1982; Xin and Bloomfield, 1999). The surround is mediated by GABAergic inhibition, it depends on activation of ON bipolar cells, and it is blocked by TTX, indicating that it arises from a spiking AC (Bloomfield and Xin, 2000) (Figure 2D-G). The NOS-1 AC satisfies all of the criteria for the mechanism generating the AII surround: it is a GABAergic [Figure 8A, B (Zhu et al., 2014)], spiking ON AC (Figure 5B, C; Figure 8C, D) that provides the majority of its synaptic output to AIIs (Figures 3 and 4; Table 2). Furthermore, NOS+ ACs are necessary for generating TTX-dependent surround inhibition in AIIs (Figure 7A, B).

Our conclusion that the NOS-1 AC provides ∼90% (Table 1) of the inhibitory input to the AII is based on the assumption—strongly supported by the anatomical evidence (Figures 3 and 4)—that a single cell type is presynaptic to all of the inhibitory synapses on the distal dendrites of AIIs. Given that every inhibitory input to the distal dendrites of three AIIs analyzed arises from a process with a stereotyped pattern of synaptic output (Figures 3C2 and 4A, B), we assume they represent a single type. The alternative seems very unlikely: that there are multiple independent populations of amacrine cell, each of which has axons identical in appearance, running in exactly the same stratum of the IPL, providing the identical pattern of 80% output to AIIs and 20% output to RBs, and matching precisely the pattern of output of the two reconstructed cells (both of which share identical patterns of inputs as well as outputs). Indeed, it is established that individual retinal neuron types are defined by their highly stereotyped patterns of connectivity (Briggman et al., 2011; Cohen and Sterling, 1990; Graydon et al., 2018; Hoggarth et al., 2015). Further, the combination of wide-field axons in the ON laminae of the IPL and narrow-field dendrites in the OFF laminae of the IPL apparently is restricted to a single population of wide-field AC (Lin and Masland, 2006; Zhu et al., 2014).

From our SBEM analysis, the multi-stratified AC presynaptic to the AII is predicted to be an ON cell that should be activated by dim scotopic stimuli owing to its input from RBs as well as from type 6 CBs, which are well-coupled to the AII network (Grimes et al., 2014b): our observations of the light responses of NOS-1 ACs (Figure 6), their synaptic connectivity (Figures 8-9), and the properties of surround suppression of GC responses to dim light stimuli (Figure 2D) collectively support the conclusion that the NOS-1 AC is the primary inhibitory neuron influencing the output of the AII network to rod-driven input. As well, it is notable that the NOS-1 AC appears to be one significant mechanism by which cone pathways can inhibit rod pathways (Lauritzen et al., 2016). And, given that the NOS-1 AC produces NO, which is thought to regulate electrical transmission between AIIs and ON cone bipolar cells (Mills and Massey, 1995), it is possible that synaptic inhibition of the AII is coupled with modulation of its electrical synapses.

### Functional relevance

When considering the stochastic nature of single photon absorption by rods, the rod integration time, and the pooling of the output of multiple rods, the ability of RBs to encode contrast in natural scenes emerges at a mean luminance of ∼10-20 R*/rod/s (Beaudoin et al., 2008). Significantly, as luminance increases from darkness to 10-20 R*/rod/s, the gain of transmission at the RBàAII synapse is reduced (Dunn et al., 2006; Dunn and Rieke, 2008), and AIIs hyperpolarize; this hyperpolarization spreads to the terminals of type 6 CBs and increases rectification at type 6àON α GC synapses to counter the influence of activity-dependent synaptic depression (Grimes et al., 2014a). Although it was suggested previously that synaptic depression at the RBàAII synapse caused the hyperpolarization of the AII (Grimes et al., 2014a), we think it more likely that synaptic inhibition is the major driver of this phenomenon because we previously demonstrated that the RBàAII synapse remains functional at backgrounds as high as 250 R*/rod/s (Ke et al., 2014).

Thus, inhibitory input from the NOS-1 AC might represent a significant modulatory mechanism within the RB pathway and could be responsible for maintaining high-fidelity signaling through a range of background luminance at which differentiating signal from noise is a particular concern. More generally, because the type 6 CB provides input to the NOS-1 AC and is coupled electrically to the AII network that is inhibited by the NOS-1 AC, we propose that a type 6 CBàNOS-1 ACàAII ACàtype 6 CB feedback circuit could maintain the rectifying nature of transmission at type 6 CB synapses by preventing excessive presynaptic depolarization under a range of lighting conditions.

## Conclusion

We uncovered the dominant inhibitory input to the AII, the central neuron in the well-studied RB pathway of the mammalian retina. The NOS-1 AC is the spiking neuron responsible for generating the TTX-sensitive, GABAergic surround that modulates AII network function across a range of lighting conditions (Bloomfield and Xin, 2000; Xin and Bloomfield, 1999). Further, our anatomical analysis also explained the ON response of a multi-stratified AC with neurites and excitatory synaptic inputs in both the ON and OFF strata of the IPL. Our study thereby extends the classification of retinal neurons that receive ON input from *en-passant* axonal synapses made by ON bipolar cells in the OFF strata of the IPL, which also includes the dopaminergic AC and the intrinsically-photosensitive ganglion cells (Dumitrescu et al., 2009; Hoshi et al., 2009; Sabbah et al., 2017). Our study demonstrates the utility of a targeted connectomic analysis coupled with neurophysiological investigation to neural circuit discovery.

## Methods

### TEM

An excised retina was fixed for one hour at room temperature with 2% glutaraldehyde in 0.15 M cacodylate buffer, washed in three changes of the same buffer, and postfixed with 1% osmium tetroxide in 0.15 M cacodylate containing 1.5% potassium ferrocyanide. A wash in three changes of distilled water followed the reduced osmium fixation and preceded an *en bloc* fix in 2% aqueous uranyl acetate. Dehydration in a graded series of ethanol (35% to 100%), and infiltration in a propylene oxide:epoxy resin series was followed by embedding and polymerization in epoxy resin. Thin sections were cut on a Reichert Ultracut E ultramicrotome, stained with 2% uranyl acetate and 0. 2% lead citrate before being viewed and photographed on a Zeiss EM10 CA transmission electron microscope.

### SBEM Analysis

Dataset k0725, a 50×210×260 µm block of fixed mouse retina imaged with voxel size 13.2×13.2×26 nm (Ding et al., 2016) was analyzed. Manual skeletonization and annotation were performed using Knossos (www.knossostool.org; (Helmstaedter et al., 2011). Tracing began at RB terminals, which were easily identified based on their size and position within the inner plexiform layer (IPL) (Mehta et al., 2014; Pallotto et al., 2015). From RB dyad synapses, AIIs were traced, and then, from sites of synaptic input, ACs were traced. All skeletons and annotations were checked by two expert observers. Voxel coordinates were tilt-corrected and normalized to the positions of the ON and OFF SACs [per (Helmstaedter et al., 2011)] identified by Ding (Ding et al., 2016). Connectivity analysis was performed using custom-written Python scripts. Skeletons were visualized in Paraview (www.paraview.org).

### Electrophysiology

All animal procedures were approved by the Institutional Animal Care and Use Committees at Yale University or the University of Maryland. Experiments used offspring of nNOS-CreER and Ai32 mice. In nNOS-CreER mice (B6; 129S-Nos1^tm1.1(cre/ERT2)/Zjh^/J; Jackson Laboratory #014541, RRID:IMSR_JAX:014541), expression of Cre recombinase is driven by endogenous Nos1 regulatory elements (Taniguchi et al., 2011), and Cre expression was induced by tamoxifen (2 mg delivered on two consecutive days) administered by either IP injection or gavage at P31 (SD = 5.6 days), and at least two weeks before the experiment. Ai32 mice (B6;129S-Gt(ROSA)26Sor^tm32(CAG-COP4*H134R/EYFP)Hze^/J; Jackson Laboratory #024109, RRID:IMSR_JAX:024109) express a Cre-dependent channelrhodopsin-2 (ChR2)/enhanced yellow fluorescent protein (eYFP) fusion protein (Madisen et al., 2012). Mice studied were heterozygous for the Cre allele and the Ai32 reporter allele.

A mouse aged between ∼2-4 months was dark adapted for one hour, and following death, the eye was enucleated and prepared for recording in Ames medium (Sigma) under infrared light using night-vision goggles connected to a dissection microscope (Park et al., 2015). In the recording chamber, the retina was perfused (∼4-6 ml/min) with warmed (31–34°C), carbogenated (95% O_2_-5% CO_2_) Ames’ medium (light response and optogenetic experiments in whole-mount retina). The retina was imaged using a custom-built two-photon fluorescence microscope controlled with ScanImage software [RRID:SCR_014307 (Borghuis et al., 2013; Borghuis et al., 2011; Pologruto et al., 2004)]. Fluorescent cells were targeted for whole-cell patch clamp recording with a Coherent Ultra II laser tuned to 910 nm (Park et al., 2015). For optogenetic experiments in retinal slices, dissection was performed in normal room light, and the retina was maintained in artificial CSF as described previously (Jarsky et al., 2010).

Electrophysiological measurements were made by whole-cell recordings with patch pipettes (tip resistance 4-11 MΩ). Membrane current or potential was amplified, digitized at 10-20 kHz, and stored (MultiClamp 700B amplifier; ITC-18 or Digidata 1440A A-D board) using either pClamp 10.0 (Molecular Devices) or IGOR Pro software (Wavemetrics). For light-evoked responses and optogenetic experiments in whole-mount retina, pipettes contained (in mM): 120 Cs-methanesulfonate, 5 TEA-Cl, 10 HEPES, 10 BAPTA, 3 NaCl, 2 QX-314-Cl, 4 ATP-Mg, 0.4 GTP-Na_2_, and 10 phosphocreatine-Tris_2_ (pH 7.3, 280 mOsm). For optogenetic experiments in retinal slices, pipettes contained (in mM): 90 Cs-methanesulfonate, 20 TEA-Cl, 1 4-AP, 10 HEPES, 1 BAPTA, 4 ATP-Mg, 0.4 GTP-Na_2_, and 8 phosphocreatine-Tris_2_.

Either Lucifer Yellow (0.05 - 0.1%) or red fluorophores (sulfarhodamine, 10 µM or Alexa 568, 60 µM) were added to the pipette solution for visualizing the cell. All drugs used for electrophysiology experiments were purchased from Tocris Biosciences, Alomone Laboratories or Sigma-Aldrich. Excitatory and inhibitory currents were recorded at holding potentials near the estimated reversal for either Cl^-^ (E_Cl_, −67 mV) or cations (E_cation_, +5 mV), after correcting for the liquid junction potential (−9 mV). Series resistance (∼10-80 MΩ) was compensated by up to 50%. Following the recording, an image of the filled cell was acquired using 2PLSM.

Light stimuli were presented using a modified video projector [peak output, 397 nm; full-width-at-half-maximum, 20 nm (Borghuis et al., 2014; Borghuis et al., 2013)] focused onto the retina through the microscope condenser. Stimuli were presented within a 4 x 3 mm area on the retina. Stimuli included contrast-reversing spots of variable diameter to measure spatial tuning (Zhang et al., 2012). For some experiments, stimuli were presented with 1-Hz temporal square-wave modulations (100% Michelson contrast) relative to a background of mean luminance that evoked ∼10^4^ photoisomerizations (R*) cone^-1^ sec^-1^ (Borghuis et al., 2014). In other experiments, stimuli were spots of dim light (4 – 40 R* rod^-1^ sec^-1^) of varying diameter presented on a dark background.

### Optogenetics

ChR2-mediated responses were recorded in the presence of drugs to block conventional photoreceptor-mediated light responses. Recordings were made in a cocktail of (in µM): L-AP4 (20); either UBP310 (50) or ACET (1-5); DNQX (50-100); and D-AP5 (50-100) (Park et al., 2015). ChR2 was activated by a high-power blue LED (ƛ_peak_, 450 or 470 nm; maximum intensity of ∼5 x 10^17^ photons s^-1^ cm^-2^) focused through the condenser onto a square (220 µm per side) area as described previously (Park et al., 2015).

### Construction and production of recombinant AAV

To generate an AAV vector backbone, we modified two plasmids procured from Addgene (#62724 and #74291). We digested plasmid #74291 with BamHI, treated with Klenow fragment, digested again with HindIII, and kept the vector backbone. The insert part of plasmid #62724 was excised by digesting with EcoRI, treated with Klenow fragment, and digested again with HindIII. The excised fragment was ligated into the vector backbone from plasmid #74291. Then, we amplified the DTA sequence by PCR, created KpnI and NheI restriction sites at each end, and subcloned the PCR products into a newly generated AAV vector. The final construct contained DTA sequence in reverse orientation surrounded by two nested pairs of incompatible loxP sites (pAAV-CAG-FLEX-NheI-DTA-KpnI-WPRE-SV40pA). The plasmid carrying DTA was obtained from Addgene (#13440).

Virus production was based on a triple-transfection, helper-free method, and the virus was purified as described previously (Byun et al., 2019; Park et al., 2015), except that we used a plasmid carrying AAV2/7m8 capsid (gift from Dr. John Flannery, University of California at Berkeley). The titer of the purified AAVs was determined by quantitative PCR using primers that recognize WPRE; the concentrated titers were >10^13^ viral genome particles/ml in all preparations. Viral stocks were stored at −80°C.

### Histology

For immunohistochemistry, animals were perfused at age 1-12 weeks. The retinas were dissected and fixed with 4% paraformaldehyde for 1 h at 4°C. For whole-mount staining, retinas were incubated with 6% donkey serum and 0.5% triton X-100 in PBS for 1 h at room temperature; and then incubated with 2% donkey serum and 0.5% triton X-100 in PBS with primary antibodies for 1–4 days at 4°C, and with secondary antibodies for 1-2 h at room temperature. For morphological analysis of recorded cells, the retina was fixed for 1 h at room temperature and reacted as described previously (Manookin et al., 2008).

Primary antibodies were used at the following concentrations: goat anti-ChAT (1:200, Millipore AB144P, RRID: AB_2079751), rabbit anti-Lucifer Yellow (1:2000, ThermoFisher Scientific A-5750, RRID: AB_2536190), rabbit anti-nNOS (1:500, ThermoFisher Scientific 61-7000, RRID: AB_2533937), guinea pig anti-nNOS (1:2000, Frontier Institute Af740, RRID: AB_2571816), rabbit anti-TH (1:1000, Millipore AB152, RRID: AB_ 390204), rabbit anti-human/rat CRF serum (1:40,000, code #PBL rC68; gift of Dr. Paul Sawchenko, Salk Institute), guinea pig anti-LHX9 (1:20,000, gift of Dr. Jane Dodd, Columbia University). Rabbit anti-nNOS was used in all cases of nNOS immunolabeling except for Figure 6F and G, in which case guinea pig anti-nNOS was used. Secondary antibodies (applied for 2 hours) were conjugated to Alexa Fluor 488, Cy3 and Cy5 (Jackson ImmunoResearch) and diluted at 1:500.

### Confocal imaging

Confocal imaging was performed using Zeiss laser scanning confocal microscopes (510, 710, or 800 models). For filled cells, a whole-mount image of the dendritic tree was acquired using a 20X air objective (NA = 0.8); in some cases, multiple images were combined as a montage. A high-resolution z-stack of the ChAT bands (i.e., cholinergic starburst AC processes, labeled by the ChAT antibody) and the filled amacrine or ganglion cell was obtained to determine their relative depth in the IPL using a 40X oil objective (NA = 1.4). Analysis of nNOS, Lhx9, CRH and TH antibody labeling was performed either with the 40X oil objective.

## Acknowledgements

Supported by EY017836 to JHS, EY014454 to JBD, EY029820 to I-JK, P30-EY026878 to Yale University, and an unrestricted grant from Research to Prevent Blindness to Yale University. We thank Tim Maugel and the Laboratory for Biological Ultrastructure (Department of Biology, UMD) for assistance with transmission electron microscopy, Alexander Baden for assistance with 3D visualization, and Kacie Furcolo and Adit Sabnis for assistance with SBEM dataset skeletonization. We thank Cole Graydon for comments on the manuscript.

